# Exploring the boundaries of microbial habitability in soil

**DOI:** 10.1101/2020.08.03.234583

**Authors:** Nicholas B. Dragone, Melisa A. Diaz, Ian Hogg, W. Berry Lyons, W. Andrew Jackson, Diana H. Wall, Byron J. Adams, Noah Fierer

## Abstract

Microbes are widely assumed to be capable of colonizing even the most challenging terrestrial surface environments on Earth given enough time. We would not expect to find surface soils uninhabited by microbes as soils typically harbor diverse microbial communities and viable microbes have been detected in soils exposed to even the most inhospitable conditions. However, if uninhabited soils do exist, we might expect to find them in Antarctica. We analyzed 204 ice-free soils collected from across a remote valley in the Transantarctic Mountains (84 – 85°S, 174 – 177°W) and were able to identify a potential limit of microbial habitability. While most of the soils we tested contained diverse microbial communities, with fungi being particularly ubiquitous, microbes could not be detected in many of the driest, higher elevation soils - results that were confirmed using cultivation-dependent, cultivation-independent, and metabolic assays. While we cannot confirm that this subset of soils is completely sterile and devoid of microbial life, our results do show that microbial habitability and activity can be restricted by near-continuous exposure to cold, dry, and salty conditions, establishing the environmental conditions that constrain habitability in terrestrial surface environments. Constant exposure to these conditions for thousands of years has generated uninhabited surface soil environments, with either no detectable microbes or conditions which are not suitable to sustain microbial activity. Such uninhabited soils are unlikely to be unique to the studied region with this work challenging expectations about where microbes might, or might not, be able to thrive on Earth and other planets.

**Significance Statement:** Certain surface soils in Antarctica have remained effectively uninhabited due to a near-continuous exposure to cold-dry-salty conditions. This is an unexpected result because soils, even those in hyper-arid deserts, typically contain detectable microorganisms. Additionally, the prevalence of fungi at the colder, drier, higher elevation sites suggests that certain fungi may in fact be better adapted than bacteria or archaea to some of the most challenging soil environments on Earth.

## Introduction

Microorganisms are nearly ubiquitous on Earth and can routinely be found in environments that include sediments of the deepest ocean trenches (1), highly acidic geothermal systems (2), and subsurface aquifers > 1 km below the Earth’s surface (3). Viable microbes have been detected in even the most inhospitable environments (4) and it is widely assumed that all environments on Earth should contain detectable microorganisms (5). This assumption is likely incorrect. Lava flows and lava lakes, for example, are inhospitable to all microbial life but only remain so for relatively short periods of time until temperatures have cooled sufficiently to permit microbial colonization (5). Few of Earth’s stable terrestrial surface environments are expected to experience conditions challenging enough to keep them uninhabited, with ‘uninhabited’ defined as environments where microbial biomass is below detection limits *or* environments in which conditions are not suitable to sustain microbial activity (5).

Uninhabited surface soils, in particular, are presumably very rare. Soils typically harbor diverse microbial communities of bacteria, fungi, and archaea (6). This holds true even for some of the most challenging soil environments found on Earth (7, 8). For example, even soils in the driest regions of the Atacama Desert contain active microbial communities despite these sites receiving ∼2 cm of precipitation annually (7). Soil microbes are impressively adept at tolerating ‘extreme’ conditions, including very high or very low temperatures, high levels of UV radiation, high salt concentrations, very low or high pH levels, and even a near-complete absence of liquid water (4, 6, 7, 9). This wide range of microbial tolerances to challenging environmental conditions suggests that, given sufficient time, all surface soils on Earth should harbor active microbial life.

Do surface soils exist where microbial life cannot be detected? If so, such ‘uninhabited’ surface soils may exist in Antarctica (8, 10). While we know that some Antarctic soils can harbor diverse and metabolically active microbial communities (11, 12), other soils harbor some of the lowest recorded levels of microbial biomass on Earth (8). Many ice-free surface soils in Antarctica are thought to have been exposed for millions of years and have remained largely unchanged over that time (13). These soils have generally been unaffected by direct anthropogenic impacts (14) and are often very isolated (15, 16). Even if microbes can be dispersed to remote soil patches via aeolian transport (16), they often face a unique combination of cold temperatures, extremely low soil water potentials, and high salt concentrations that restrict the activity and survival of all but a few specifically adapted taxa (4, 8). It seems likely that some isolated Antarctic soils may have remained effectively uninhabited by microbes due to these constraints.

To explore the limits of microbial habitability in terrestrial soil environments, we sampled Antarctic surface soils from 10 distinct features along the Shackleton Glacier in the Transantarctic Mountains (84 – 85°S, 174 – 177°W). We used a suite of microbiological approaches, including cultivation-dependent, cultivation-independent, and metabolic assays, to answer two questions: Can we find uninhabited soils in Antarctica with no detectable signs of microbial life? and, if so, What environmental conditions might be limiting the activity of microorganisms in these inland soils?

## Results

### Microbial communities of the Shackleton Glacier Region

The 204 individual soils collected for this study represent a broad range of ice-free sites found across the Shackleton Glacier region. Sites ranged in elevation from ∼150 m to 2221 m above sea level (m.a.s.l) and most soils are likely to have been exposed for prolonged periods of time, with the approximate time since last wetting (estimated from ClO_4_^-^ concentrations, 17,18) ranging from <10 years to >2 million years (mean: ∼20,000 years). Not surprisingly given the absence of plants in this region, the measured soil organic carbon concentrations were low, from ∼3 to 60 mg·g soil^-1^ (mean = 13 mg g soil^-1^), and most of the collected soils have high soluble salt concentrations (Figure 3, Figure S5). As expected, the soils contained almost no water at the time of collection (0.001 to 0.11 g H_2_O·g dry soil^-1^, mean of 0.02 g H_2_O·g dry soil^-1^). In general, soils located further inland at higher elevations were drier, saltier, and contained less organic carbon (19). These soil characteristics are not unique to the Shackleton Glacier region as ice-free soils found in other regions of Antarctica have comparable edaphic characteristics (15,20).

**Figure 1.**
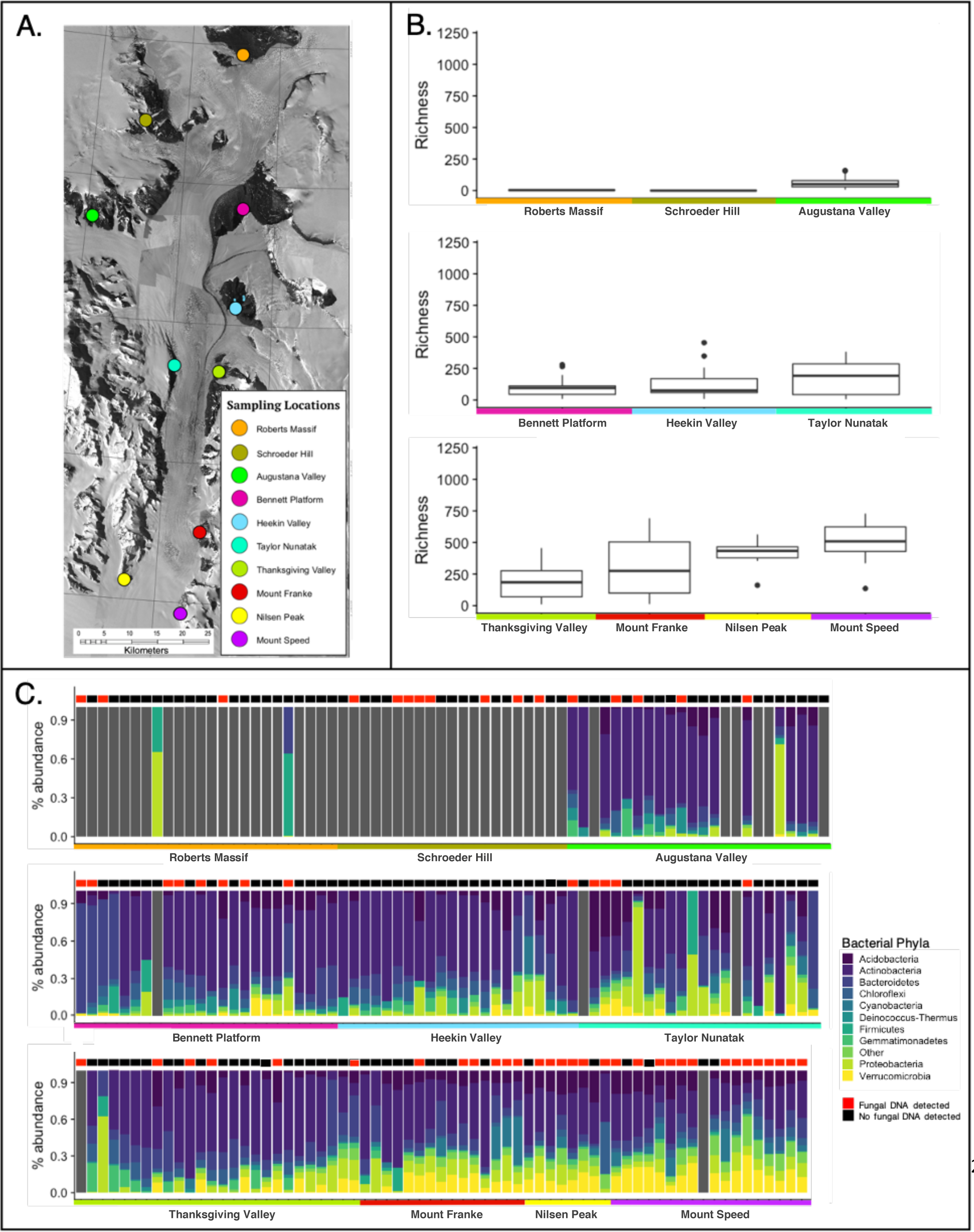
Richness and composition of the soil bacterial communities across the Shackleton Glacier region. (A) A map of the Shackleton Glacier region (Lat: 84 - 85°S, Long: 174 – 177°W) with the locations of the 10 features where samples were collected indicated by the colored dots. (B) The richness of the bacterial communities (number of bacterial phylotypes detected per sample) from each of the 10 sampled features. Box and whisker plots indicate the distribution of bacterial richness levels (number of phylotypes per sample) across all samples from each individual feature. (C) Relative abundances of the dominant bacterial phyla from the 204 surface soil samples analyzed. Samples are grouped by feature and organized top to bottom from high elevation to low elevation (higher elevation sites are further south). Samples with no amplifiable bacterial DNA are represented by grey bars. The presence/absence of amplifiable fungal DNA is indicated by the red and black boxes over each bar.

**Figure 2.**
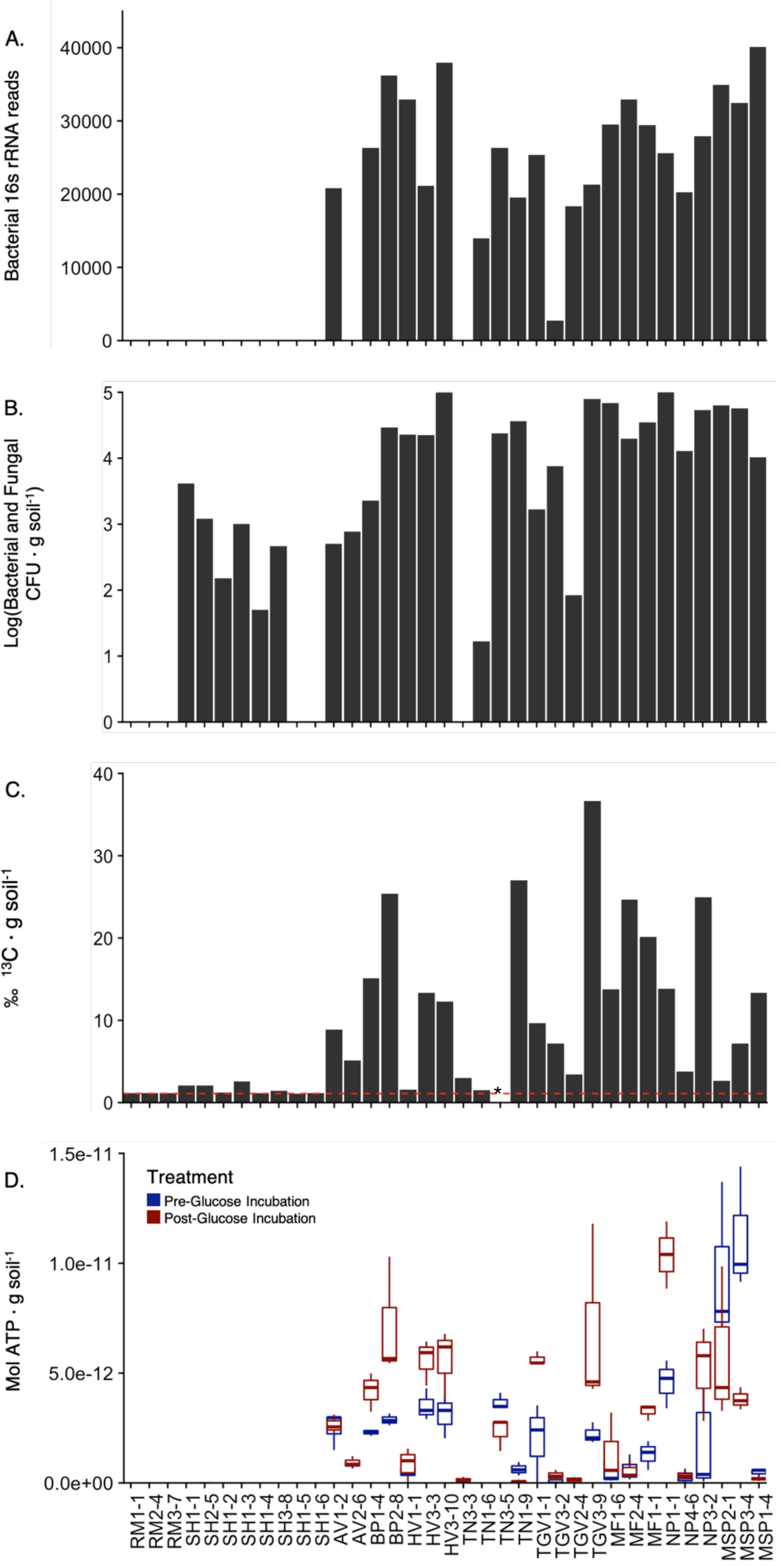
The four tests used to identify detect microorganisms and/or microbial activity in soils. (A) The number of bacterial 16S rRNA reads that were classified to at least the phylum level of resolution for the subset of samples (n=35) used to verify the results of the culture-independent sequencing. Samples with a read number less than the threshold determined to identify and reliably detect prokaryotic DNA (see Methods) have been omitted. For all panels, samples are arranged on the x-axis by elevation (higher elevation features on the left). (B) The log transformed total colony counts (Log(CFU·g soil^-1^)) of microorganisms (both bacteria and fungi) grown from each of the 35 samples. The reported total cell numbers are the cumulative number of colonies that grew on each plate over the three-month incubation period. Samples with no bar had no growth, i.e. no colonies detected. (C) The ^13^C ‰ released as CO_2_ from the glucose-amended soils as calculated from the unautoclaved replicates of each of the 35 samples. The dashed red line indicates the ^13^C ‰ measured in the blank samples. 3 samples (RM3-7, SH1-5, SH1-6) had ^13^C ‰ from their unautoclaved replicates that was not significantly different from the values calculated from either their autoclaved replicated or the blank tubes and were below detection limits. Sample TN3-5, indicated by a star, was not successfully measured. (D) ATP concentrations measured from the soils. Blue boxes were baseline readings performed on the soils before any glucose was added and before any incubation. Red boxes are from paired soil sub-samples amended with glucose and incubated for 24 h. Samples that are blank indicate that the ATP readings were below detection limits.

**Figure 3.**
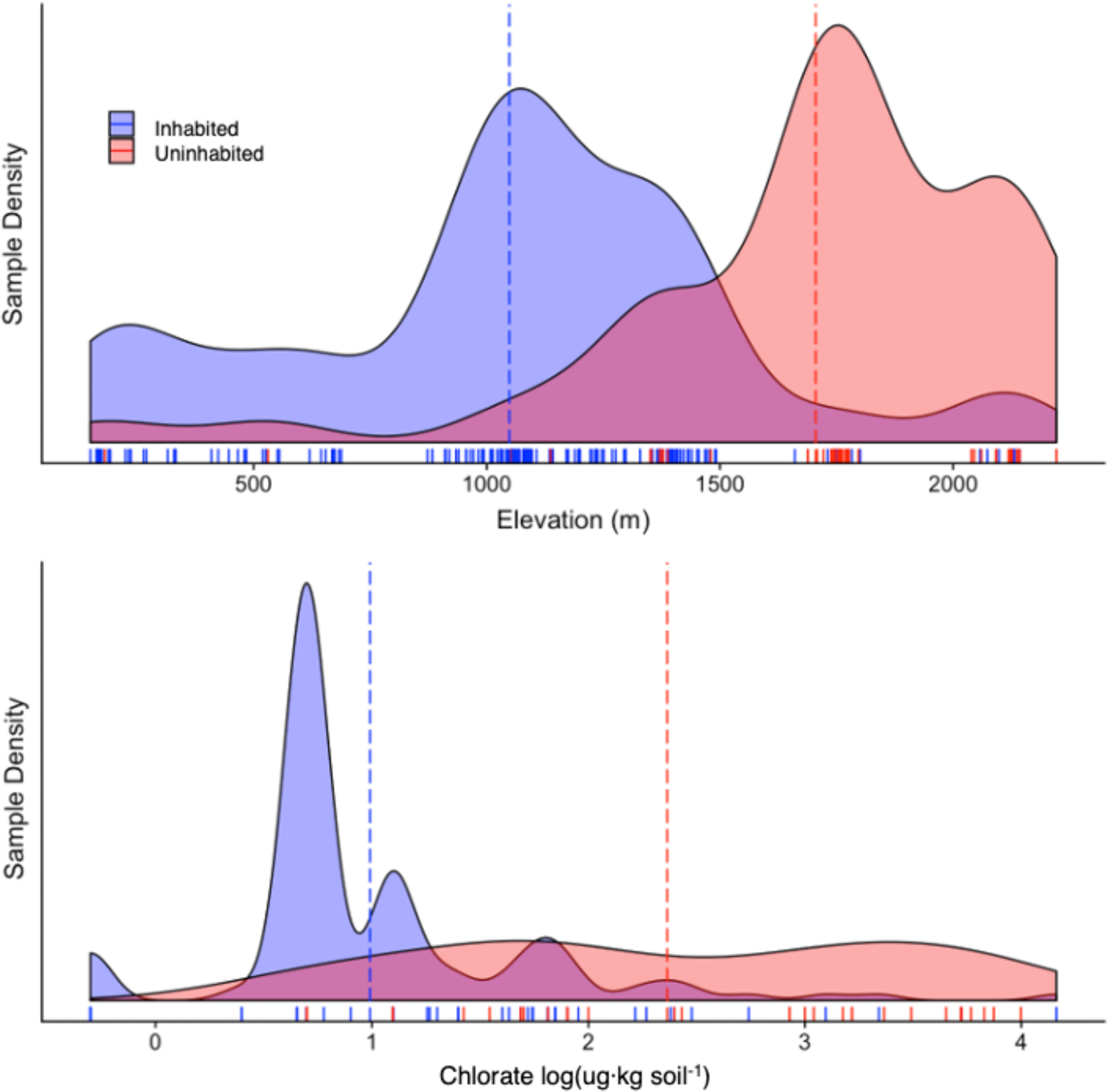
The best predictors of the habitability of a soil are elevation and chlorate concentration based on the random forest model. The distribution of inhabited and uninhabited soils across the Shackleton Glacier Region in relationship to elevation (A) and soil chlorate concentrations (B). Y values show frequency density of each group with n=204 for elevation and n= 169 for chlorate. The area under each curve is equal to 1. Concentrations of chlorate ions (µg ·kg soil^-1^) have been log transformed. Dashed lines indicate the mean value for each group. To see the distribution of all other variables tested in the random forest model and the model results, see Figure S5 and Table S4.

Of the 204 soil samples analyzed, 80% of the samples yielded sufficient amounts of PCR-amplifiable DNA to characterize the bacterial and/or fungal taxa found in these soils using cultivation-independent marker gene sequencing. As expected, the number of taxa identified per soil sample (microbial richness) was low with a mean of 205 bacterial phylotypes (2 – 729 exact sequence variants, ESVs) and only 5.5 fungal phylotypes (1 – 45 ESVs) detected per sample. In general, the soils from higher elevation features farther away from the Ross Ice Shelf had less diverse fungal and bacterial communities (Figure 1, Figure S1, Data S1, Data S2).

The bacterial phyla with the highest relative abundances across all soils were: Bacteroidetes, Proteobacteria, Actinobacteria, and Acidobacteria, with the dominant taxa members of the families *Solirubrobacteraceae, Blastocatellaceae, Chitinophagaceae*, and *Rubrobacteriaceae* (Figure 1C, Data S1). The composition of the bacterial communities in these soils is consistent with results obtained using similar cultivation-independent analyses of other Antarctic soils, including those found in the McMurdo Dry Valley region (11, 21, 22). The fungal communities in these soils were dominated by members of the Ascomycota and Basidiomycota phyla, including members of the families *Herpotrichiellaceae, Trapeliaceae, Verrucariaceae, Filobasidiaceae, Mortierellaceae, Stereocaulaceae* (Figure S1, Data S2). While there have been relatively few comparable studies of fungi in Antarctic soils, the dominant taxa identified are generally similar to those found in other Antarctic soils (23, 24). Archaeal sequences associated with the phyla Thaumarchaeota, family Nitrososphaeraceae, were detected in 60 samples, but these archaeal sequences made up a maximum of 1.5% of all the 16S rRNA gene reads per sample (mean = 0.08% of reads per sample, Data S1).

We found that 20% of the soils (40 out of 204 soils) had no amplifiable microbial DNA as determined by our cultivation-independent sequencing approach (see methods for more details). This is an unusual result for a culture-independent analysis of soil as the microbial communities found in a wide range of soils across the globe have been successfully characterized using similar techniques (25), and we have shown that the soils themselves do not appear to be inhibiting DNA extraction or PCR amplification (Figure S4). We recognize that these results do not prove that these soils are completely sterile as confirming the absence of microbial life is difficult (5) and we could have further optimized our methods to try to detect microbial DNA (e.g. extracting DNA from larger sample volumes, testing a range of different extraction kits and PCR protocols). Instead, we note that while all of these soils have very low biomass, 20% of the soils failed to yield detectable amounts of amplifiable bacterial, archaeal, or fungal DNA using techniques that are routinely used to successfully characterize microbial communities in soils from different regions of Antarctica (8, 22, 26).

### Habitability Tests

To test whether we can detect microbial life and activity in soils where microbial DNA was below detectable levels (using cultivation-independent sequencing), we analyzed a subset of 35 samples with culture-based methods, metabolic assays to detect ^13^C-glucose mineralization, and ATP quantification (in soils amended with and without glucose). This subset of 35 samples included both soils with and without microbial DNA (as determined by the cultivation-independent sequencing, Figure 1A, Figure S3) with these soils collected from transects on each of the 10 features and spanning a range of edaphic properties (Table S2).

The 35 soil samples were plated on 17 different types of solid culture media. All media targeted aerobic microorganisms (both heterotrophs and autotrophs), as obligate anaerobes are unlikely to be present in these extremely dry surface soils. This cultivation effort supports the results from the cultivation-independent analyses (Figure 2B). Out of the 35 samples tested, 4 samples did not grow any colonies on any of the different media types after three months. In general, those samples which did not yield any detectable microbial DNA also failed to yield any colonies, even when using such a broad range of aerobic media and relatively long incubation times. We acknowledge that our cultivation-based survey will not have captured all taxa potentially living in these samples as alternative cultivation strategies could always be employed, but the extensive cultivation-based analyses broadly agree with our cultivation-independent analyses in demonstrating that microbes are undetectable in an appreciable subset of the soil samples.

Of those soils (29 out of 35) which yielded visible bacterial or fungal colonies after 8 weeks, estimated cell numbers were highly variable, ranging from <20 to >130,000 CFU · g soil^-1^ (Figure 2C, Data S3). Even the soil with the highest cultivable cell numbers had values that were 50 times smaller than the ‘positive control’ soil collected from a lawn in Colorado (7.4 x 10^6^ CFU · g soil^-1^). These results are not surprising as we would expect ice-free Antarctic soils to have relatively low cultivable cell numbers (22). However, it is worth noting that, while bacterial isolates dominated the culture collection, we were also able to culture and isolate viable fungi from 14 of these soils including some where no bacteria could be cultured (Figure 2B). These results extend the cultivation-independent fungal analyses highlighted above in demonstrating that even viable fungi are often present and that bacteria are not the sole members of these Antarctic soil microbial communities. This prevalence of fungi in soils from higher elevation sites is consistent with the culture-independent sequencing results (Figure 1B, Figure S2) and suggests that certain fungi may in fact be better adapted than bacteria or archaea to some of the most challenging soil environments on Earth.

To further assess microbial activity, we performed an assay to detect microbial activity in these soils based on the catabolic mineralization of ^13^C-labeled glucose added to soils (Figure 2C, Data S3). We found that the 32 of the 35 samples tested had ^13^C-CO_2_ values that were significantly higher than their autoclaved replicates and the corresponding control blanks. Three of the 35 samples had non-autoclaved ^13^C-CO_2_ values that were indistinguishable from their paired autoclaved replicates or from the soil-free blanks. This suggests that we were unable to detect glucose mineralization resulting from microbial activity in these 3 soil samples (Table S3). This same subset of soils also had no culturable bacteria or fungi (Figure 2B), confirming that microbial cells were not detectable using the methods employed.

As a final test for microbial activity, we measured soil ATP concentrations both with and without glucose amendments (Figure 2D, Data S3). In theory, ATP assays should be able to detect very low levels of microbial activity that might not be detectable using other methods (26). We found that 21 of 35 of the soils had measurable ATP concentrations with the addition of the PBS buffer only, while 23 of 35 had measurable ATP activity after amendment with glucose and incubated for 24 h. ATP production increased after glucose amendment in nearly all of the samples that had measurable ATP in the corresponding unamended soil. All reagent blanks and blank wells were below ATP detection limits of 10^−15^ mol ATP ·g soil^-1^. ATP concentrations in 12 of the 35 samples were below detection both with and without glucose amendment. These results suggest that there was no measurable microbial ATP in these soils, further evidence for a lack of actual or potential microbial activity in a subset of the samples collected.

Together these four distinct methods (cultivation-independent marker gene sequencing, the extensive cultivation effort, the whole-soil ^13^C-glucose metabolic assay, and the ATP assays, Figure 2) yielded similar results. We were unable to detect microorganisms or microbial activity in a subset of the samples regardless of the methods employed: no colonies grew on any aerobic media type, no microbial metabolic activity was detected, and we could not detect any measurable ATP production. Although microscopy-based cell counting can be used to quantify microbial cell numbers in higher biomass soils, we did not use such an approach here given the problems associated with distinguishing between cells and soil particles in these soil types (8, 28, 29).

We recognize that proving the absence of living microbes is not easy as there are always more strategies that could be employed. However, our use of multiple distinct methods suggests that any microbial life that may exist in these soils is below the limit of detection of tests that are routinely used in other low biomass microbial systems (2, 7, 25, 30, 31, 32). We recognize the possibility that some of these soils may contain low levels of microbial biomass that we were unable to detect with these analytical approaches which is why we do not conclude that the soils are sterile. Rather we emphasize that microbial biomass is sufficiently low in a subset of samples to be undetectable, i.e. a subset of these surface soils collected from Antarctica are effectively uninhabited.

### Why are some soils uninhabited?

We next sought to determine what soil or site characteristics differentiated those soils that were apparently uninhabited from those in the same region that had detectable microbial activity. We found that elevation and soil chlorate concentrations were the best predictors of whether a soil was uninhabited or inhabited (Figure 3, Figure S5, Table S4). Soils from lower elevations were more likely to have detectable microorganisms (37% of soils from sites > 1190 m.a.s.l were uninhabited while only 3% of soils from sites <1190 m.a.s.l. were uninhabited) and soils with lower chlorate concentrations were more likely to have detectable microorganisms (uninhabited and inhabited soils had average chlorate concentrations of 1675± 2595 µg·kg^-1^ and 161 ±1273 µg·kg^-1^, respectively). Uninhabited soils also had higher median total salt and perchlorate concentrations (Figure S5), but these differences were not significant (Table S4).

Elevation and chlorate concentration may be the best predictors of whether a soil is uninhabited or contains active microorganisms, but it is unlikely that these factors, singly or in combination, are restricting organisms from being active in these soils. We know, for example, that there are soils at much higher elevations that have active microbial communities (7, 32) and microorganisms have been found in soils with higher concentrations of deposited chlorate in the Atacama Desert and in the McMurdo Dry Valleys (33, 34). Instead, a more parsimonious explanation is that these variables are indicative of a suite of environmental properties that together may have kept these soils free of detectable microbial activity. Elevation likely correlates with a number of variables that increase in magnitude further inland and higher in elevation in the Shackleton Glacier region that may reduce microbial viability: higher UV radiation, lower temperatures, lower water availability, and increased soluble salt concentrations (19). The accumulation of high concentrations of chlorate salts, on the other hand, may influence habitability due to its toxic properties but this seems unlikely due to the proximity of inhabited soils to uninhabited soils across this region (Figure 1). Instead, high concentrations of chlorate, a highly soluble molecule, may indicate that these soils seldom, if ever, have had sufficient amounts of liquid water to sustain active microbial communities (34).

There is not likely a single environmental factor that causes some of these Antarctic soils to be uninhabited. Instead, it is likely that the constant exposure to deposited salts, low temperatures, and extremely low water potentials over thousands of years have created surface environmental conditions that repress the effective colonization, activity, and establishment of those microbes that may reach these sites via aeolian transport (16). Those inhabited soils located in close proximity to uninhabited soils may harbor microbes due to transient fluctuations in temperature, water availability, and salt concentrations. Clearly microbial activity in challenging soil environments is not ubiquitously distributed and local, landscape-level differences in environmental conditions can impose limits on microbial habitability in certain soils.

## Discussion

Our finding that some ice-free soils in the Shackleton Glacier region of Antarctica are uninhabited is significant. These soils were neither recently formed, nor did they come from completely isolated areas. Instead, these soils are likely very old, and some were found on features in reasonably close proximity (within 50 m) to soils that contained detectable microbial life. This distinction is noteworthy as young soils may not have had a sufficient amount of time for microbial colonization. Instead it appears that a unique set of environmental conditions, those associated with higher elevation sites further inland and a near complete lack of liquid water over time, create conditions unfavorable to microbial activity. Soils typically contain large numbers of active microorganisms (often >1000 kg of microbial biomass carbon per hectare, 4) and even dry polar desert soils from the McMurdo Dry Valleys tend to have detectable, diverse, and active microbial communities (11, 35). To find any uninhabited soils is unexpected.

Ice-free soils are not unique to the Shackleton Glacier region. More than 45,000 km^2^ of ice-free terrestrial surfaces can be found in Antarctica >5km from the coast (36). The environmental and geochemical characteristics acting on these inland, high elevation, ice-free soils are similar to those we observed in the Shackleton Glacier region (15, 20) which suggests that uninhabited patches lacking active microorganisms may be common elsewhere in Antarctica. This possibility is supported by the research of Goordial et al. (2016), whose work on soils in the University Valley also suggests that the unique combination of conditions found in Antarctica can severely limit microbial activity and survival (8). A similar constraint on habitability may also exist at higher elevation sites in the Altiplano of South America (7), but the absence of active microbial life in these soils remains hypothetical. Regardless, uninhabited soils are likely very rare outside of Antarctica, as other ultra-arid regions, like the Atacama and Namib Deserts, harbor detectable microbial communities (37).

We hypothesize that these soils may not remain uninhabited for long. Antarctic terrestrial systems have changed very little since the Last Glacial Maximum due to relatively stable climatic conditions (38, 39). Increases in temperature, precipitation and moisture availability are predicted to accelerate at high elevations in coming years due to ongoing climate change (35). These anticipated environmental changes would likely expand the range of habitable soil environments in Antarctica (40) and decrease the likelihood of finding uninhabited soils in the future.

Finally, the existence of these uninhabited soils has important implications for the ongoing search for life on Mars. The combination of conditions found in the surface soils of the Shackleton Glacier region (low temperatures, low moisture availability, high concentrations of salt) are similar to those found on the surface of Mars (5, 41, 42). Given that Martian soils are much older, experience similar or even harsher conditions, and contain even higher concentrations of the same salts (including perchlorate and chlorate salts, 41, 42), our results suggest that searching for active life in surface soils on Mars is unlikely to return positive results.

## Conclusions

The existence of uninhabited Antarctic soils, those lacking detectable microbial life, is significant to the study of terrestrial biology and astrobiology. We are not suggesting that we have found “lifeless” or “sterile” soils, nor have we identified the low temperature threshold for life (many locations on Earth maintain lower temperatures than the Shackleton Glacier and contain active microorganisms, 4). However, our inability to detect microbes or microbial activity in certain soils suggests that these surface soils represent a limit to microbial activity and survival driven by the cold, dry, and salty environmental conditions. This phenomenon is supported by previous work performed elsewhere in Antarctica (8) and is a different type of “limit to life” than what might be found in a hot, acidic environment (4, 43). Acknowledging that certain Antarctic soils may be uninhabited will allow us to better understand the adaptations that allow organisms to survive and remain active in these unique, challenging environments and predict what other soils on Earth may be similarly uninhabited. Finally, understanding patterns of terrestrial habitability on our planet will set the groundwork to better predict where microbes might, or might not, be found on other planets.

## Materials and Methods

### Sample Collection

Soil samples were collected from the Shackleton Glacier region from December 2017-January 2018. A total of 204 soils were collected from ten different features running the length of the valley. These features represent a range of elevations (150 - 2221 m) across a 120 km north-south distance spanning from the Ross Ice Shelf to the Polar Plateau (Figure 1). Between 14 and 26 soil samples were collected along elevational transects located on each of ten features to maximize variation in soil characteristics and soil exposure times (amount of time at the surface and uncovered by glacial ice) at each feature. Soils (0 - 5 cm depth) were collected in sterile polyethylene bags using clean hand trowels. GPS coordinates, photographs of the soil surface, elevation, and other metadata were collected at the time of soil sample collection. All soils were transported to the field camp in insulated coolers where they were frozen at −20°C and remained frozen until processed at the University of Colorado in Boulder, Colorado, USA.

### DNA extractions

DNA was extracted from all of the collected samples in a laminar flow hood. After mixing 1 g of each soil in 1 mL of sterile PCR-grade water, DNA was extracted from a 500 µl aliquot of the soil slurry using the Qiagen DNeasy^®^ Powersoil^®^ HTP 96 Kit (Qiagen, Germantown, MD, USA) following the manufacturer’s recommendations. A total of 6 extraction blanks (2 per 96-well plate) were included to test for any possible contamination introduced during DNA extraction.

### Cultivation-independent microbial analyses via marker gene sequencing

The DNA aliquots extracted from each of the 204 soils and the 6 extraction blanks, were PCR-amplified using a primer pair that targets the hypervariable V4 region of the archaeal and bacterial 16S rRNA gene (515F: 5’-GTGCCAGCMGCCGCGGTAA-3’ and 806-R: 5′-GGACTACHVGGGTWTCTAAT-3′). To assess the fungal communities that may be found in these soils, we also conducted a separate set of PCR amplifications using a primer pair (ITS1-F: 5’-CTTGGTCATTTAGAGGAAGTAA-3’ and ITS2-R: 5′-GCTGCGTTCTTCATCGATGC-3′) that target the internal transcribed spacer of the fungal ribosomal RNA (rRNA) operon. Three no template PCR blanks (1 per 96 well plate) were run for each set of amplifications. Both primer sets included the appropriate Illumina adapters and unique 12 - bp barcode sequences to permit multiplexed sequencing (44). PCR was performed using GoTaq^®^ Hot Start PCR Master Mix (Promega, Madison, WI, USA) in 25 µL reaction volumes. Cycling parameters for both primer sets consisted of an initial denaturation step at 94 °C for 3 min, followed by 35 cycles of denaturation at 94 °C (45 s), annealing at 50 °C (60 s), extension at 70 °C (90 s), and a final extension step at 72 °C for 10 min. Both sets of amplified products were cleaned and normalized to equimolar concentrations using SequalPrep™ Normalization Plates (Thermo Fisher Scientific, Carlsbad, CA, USA) with the 16S and ITS rRNA gene amplicons sequenced on separate Illumina MiSeq runs (Illumina, San Diego, CA, USA) using the V2 2 x 150 bp and 2 x 250 bp paired-end Illumina sequencing kits (for the 16S rRNA gene and ITS amplicons, respectively).

16S rRNA gene sequences were processed using the DADA2 pipeline (45). Sequences were quality filtered and clustered into exact sequence variants (ESVs). Taxonomic information was assigned to ESVs using a naive Bayesian classifier method (46), which takes the set of ESVs generated and compares them to a training set of reference sequences from the 16S rRNA bacterial and archaeal SILVA database (47, 48). A minimum bootstrapping threshold required to return a taxonomic classification of 50% similarity was used for analysis. ESVs associated with chloroplast, mitochondria, eukaryotes, and those unassigned to the phylum level (477 ESVs) were removed prior to downstream analyses. Extraction blanks yielded an average of 420 reads (0 – 1831 reads) and these reads came from 14 phylotypes from 11 families *Rubrobacteriaceae, Sphingomonadaceae, Sulfruspirillaceae, Xanthobacteraceae, Burkholderiaceae, Sporolactobacillus, Clostridiaceae, Bacillaceae, Planococcaceae, Deinococcaceae, Enterobacteriaceae*, some of which have been classified as bacterial taxa commonly associated with reagent contamination (49). These 14 ESVs found in the extraction blanks represented 100% of the reads from these blanks, while these 14 ESVs only accounted for 0.31% of the reads from all extracted soil samples on average. No-template PCR blanks yielded 311 reads on average (126 - 586) and ESVs associated with these reads were common in other samples (including taxa within the families: *Blastocatellaceae, Soilrubrobacteriaceae, Clostridiaceae, Paenibacillaceae, Chitinophagaceae*, and *Rubrobacteriaceae*) and these taxa are not typical reagent contaminants (49), but are instead most likely derived from the soil samples and represent ‘tag switching’ events (50). For this reason, we used 586 reads per sample as a threshold for determining that we could reliably detect prokaryotic DNA using this DNA sequencing approach. All soils with <586 reads were considered to have no detectable PCR-amplifiable prokaryotic 16S rRNA genes. Across the 153 samples that had a sufficient number of bacterial or archaeal 16S rRNA gene reads for downstream analyses, the mean number of reads per sample was 25912 (629 - 58150) while the mean number of reads in the 51 samples that did not meet the threshold determined from the analysis of the ‘blank’ samples was only 125 reads per sample.

Fungal ITS sequences were also processed using the DADA2 pipeline (42). Sequences were quality filtered and clustered into exact sequence variants (ESV). Taxonomic information was assigned to ESVs using the same naive Bayesian classifier method described previously^43^ but with the UNITE database (51). A minimum bootstrapping threshold required to return a taxonomic classification was set at 85% similarity (51, 52). All 6 of the extraction blanks and 3 no template PCR controls had zero reads after processing. We used all ESVs that could be classified to at least a fungal phylum for our analysis. After this filtering, 143 of the 204 samples had no remaining ITS reads. These 143 samples with no identifiable fungal ITS sequences were considered to have no amplifiable fungal DNA. The remaining 61 samples had a mean of 5886 reads per sample and were considered to have a sufficient number of fungal ITS reads for downstream analyses.

### DNA extraction and PCR inhibition test

From the cultivation-independent ITS and 16S rRNA gene sequencing analyses described above, we identified ∼ 20% of samples (40 out of the 204 individual soils) that had no amplifiable DNA from the targeted bacteria, archaea, or fungal marker genes. To confirm that the PCR amplification of microbial DNA from these 40 samples was not simply a product of DNA extraction problems or PCR inhibition, we performed an inhibition test. For this test, we used two 250 mg subsamples of three soils that did not PCR amplify and one soil that successfully amplified. Following the methods described by Warren-Rhodes et al. (2019), one sub-sample of each of the four soils was extracted after adding a 50 µL suspension of *E. coli* in trypticase soy broth at a concentration of 10^7^ CFU (53). The other subsample was extracted with no addition of *E. coli*, with the DNA extractions on all samples conducted as described above. Extracted DNA was amplified using the 16S rRNA gene-targeting primers, reagents, and PCR conditions described above. The concentrations of both the extracted gDNA and amplified product from each soil sample replicate were measured with a Molecular Devices SpectraMax M2 microplate reader (Molecular Devices, LLC, San Jose, CA, USA) using an Invitrogen Quant-iT™ Picogreen™ dsDNA Assay Kit (Thermo Fisher Scientific, Carlsbad, CA, USA). Amplified products were also visualized on a 2% agarose gel. No signs of inhibition were detected, i.e. the DNA from all sub-samples amended with *E. coli* cells were successfully PCR-amplified (Supp. Figure 4). Thus, we can conclude that our failure to recover 16S and ITS rRNA gene reads from these 44 samples was not likely a result of PCR inhibition, but rather any microbial DNA in these samples was simply not detectable using our cultivation-independent sequencing approach.

### Cultivation-dependent analyses

As an added test of the presence of microbial cells, we attempted to cultivate microbes from a subset of 35 soil samples (Supp. Table 2).These 35 soils were selected to represent the range of environmental gradients encompassed by the larger dataset and included samples from all 10 features within the Shackleton Glacier region, including both ‘inhabited’ and ‘uninhabited’ soils (as determined by the cultivation-independent sequencing). These 35 soil samples were plated on 17 different types of media (Table S1, 54 – 59) to capture a broad diversity of microbial taxa that could be living in these soils. One gram of each soil sample was homogenized with 1 mL of water and 60 µL of the resulting soil slurry was pipetted onto each of the 58 cm^2^ plates and spread across the plates using flame sterilized cell spreaders. ‘Blank’ plates inoculated with only 60 µL sterile water were prepared for each media type and handled in an identical manner to the 595 plates inoculated with the soil slurries. One ‘positive control’ soil sample was collected from a lawn on the University of Colorado Boulder’s main campus and was plated on all media types using the same techniques. The TSA 24 plates, the PH, and PA plates were all left to incubate at 24°C while the rest of the plates were incubated at 4°C. All plates, besides the PA and PH plates, were left to incubate in the dark. Plated samples, and the uninoculated control plates, remained under these conditions for 8 weeks. Photos of the plates were taken each week. Every distinct colony on each of the 629 petri plates was counted weekly from these photos and microbial cell numbers (CFU · g soil^-1^) were calculated from these colony counts. The reported total cell numbers are the cumulative number of colonies that grew on each plate over the 8-week incubation period. At the end of the 8-week incubation, those plates that had no visible colonies (307 of 629 plates) were incubated for an additional month. No colonies grew on any of the blank plates over the three-month incubation.

### ^13^C glucose metabolism assay

To further verify whether there were, indeed, some soil samples with no detectable microbes and to complement the cultivation-independent and dependent approaches described above, we conducted a ^13^C - glucose metabolic assay with the subset of 35 soil samples described above. We used ^13^C-glucose as a substrate because glucose should be readily catabolized by most heterotrophic microbes. Two 1 g replicate sub-samples of each of the 35 soils were placed in sterile glass tubes. Three soil samples (including the one used in the culturing experiment) collected from the University of Colorado Boulder’s main campus were prepared in the same way as the Shackleton Glacier samples to serve as ‘positive’ controls. One of the replicate sub-samples from each of the 38 soil samples, plus 6 soil-free blank tubes, was autoclaved on a gravity cycle with a 60-minute sterilization and a 30-minute drying cycle so we could assess metabolic activity in paired autoclaved versus un-autoclaved sub-samples of each soil. A 250 µL solution ^13^C - glucose (99 atom% U-^13^C, Cambridge Isotope Laboratories, Tewksbury, MA, USA) dissolved in H_2_O was pipetted directly into each soil, an addition of approximately 250 µg glucose C per gram soil. This amount of glucose added to the soils is comparable to the glucose amendments used in other comparable soil metabolic assays (50, 61). A 2 mL cryotube containing 1 mL of 1N sodium hydroxide solution was placed in each tube before all tubes were sealed with an airtight cap. We prepared a total of 76 soil incubations (35 Antarctic soils + 3 Colorado soils with and without soil autoclaving) and 6 blanks (3 tubes with NaOH traps without soil or glucose, and 3 tubes without NaOH traps without soil but with glucose). All tubes were incubated for one month at 4°C after which an aliquot of the NaOH trap from each tube was transferred to pre-evacuated glass vials filled with helium gas. The NaOH traps were acidified with 1 mL of concentrated phosphoric acid to release CO_2_ into the evacuated headspace. The quantity and isotopic composition of the released ^13^CO_2_ was determined using a ThermoFisher Scientific Gasbench II (Thermo Fisher Scientific, Waltham, MA, USA) coupled to a Delta V Isotope Ratio Mass Spectrometer (Thermo Fisher Scientific, Waltham, MA, USA). Data from the mass spectrometer was processed using the R package ‘isoprocessCUBES’. Sample readings were corrected using the IAEA reference standard NBS18 (62). The delta ^13^C values were converted to fraction of ^13^C per mil, expressed as ‰, using the international standards of V-PDB (Vienna Pee Dee Belemnite, 63). Positive glucose mineralization was detected in all measurements, including in control blanks who had an average reading of 1.095 ‰ ^13^C · g soil^**-1**^. Samples were identified as harboring active microbes based on comparisons between the replicate unautoclaved and autoclaved samples and positive controls. Significance was calculated using Kruskal-Wallis tests as implemented in R with differences between groups determined by a Nemenyi test calculated using the kruskalmc function in the R package ‘pgirmess.’

### ATP assay

To further confirm the results of the ^13^C glucose metabolism assay, we conducted an ATP assay with the same subset of 35 soils to test for any bacterial activity. One-gram subsamples of each of the soils were placed in sterile 2ml microcentrifuge tubes. Tubes were homogenized into a soil slurry with 1 mL of PBS. Three 100 µl aliquots of each homogenized soil slurry (35 Antarctic soils + 3 Colorado soils) were pipetted into black, 96-well Costar Assay plates (Corning Inc., Corning, NY, USA) and ATP production was analyzed using the Promega BacTiter-Glo™ Microbial Cell Viability Assay (Promega, Madison, WI, USA) following the manufacturer’s instructions. This assay has been used in other studies to detect low levels of bacterial activity on simulated Martian environments (64). Luminescence readings were performed using a Biotek Synergy™ HT Plate Reader (Biotek Instruments, Inc., Winooski, VT, USA). Six reagent “blanks” (3 wells containing just the sterile PCR-grade water used to make the soil slurries, 3 wells with water and glucose), and 6 blank wells were measured using the same methods. After this “baseline” reading, a 250 µl filter-sterilized solution of glucose dissolved in H_2_O was added to each of the soil slurries, an addition of approximately 250 µg glucose C per gram soil. Soils were mixed and were then left to incubate at 4°C. After a 24-hour incubation the soil slurries were measured again following the previously described methods, with the same reagent blanks and blank wells.

ATP concentrations were calculated from luminescence readings based on a standard curve (R^2^=0.99) generated from a 10-fold serial dilution (10^−10^ - 10^−15^ mols) of a Tris-buffered ATP standard (Thermo Fisher Scientific, Hampton, NH, USA). Based on this standard curve, we determine that we were effectively able to detect ATP concentrations >10^−15^ mol ATP ·g soil^-1^.

### Soil Geochemical Analyses

To determine soil water content, 50 grams of soil were put into aluminum weigh dishes and dried at 105°C for 24 hrs. % H_2_O was determined by the difference between the dried and initial weights of the soils. Since studies have shown that organic matter can and does degrade at this temperature, the water content analysis was done on a separate aliquot of samples.

Soils were leached at a 1:5, soil to DI water ratio and filtered through 0.45µm nucleopore filters. Anions were analyzed using a Dionex ICS-2100 ion chromatograph and an AS-DV automated sampler. Cations were analyzed by optical emission spectrum on a PerkinElmer Optima 8300 inductively coupled plasma mass spectrometer (ICP-OES)(19). Relative soil exposure ages were estimated using perchlorate concentrations in 1:5 soil to water leaches and annual fluxes from the McMurdo Dry Valleys, Antarctica. Fluxes are estimated to range from 1 to 4 µg m^-2^ yr^-1^ and average 2 µg m^-2^ yr^-1^ (17). Perchlorate salts are highly soluble in water and the primary source is wet and dry atmospheric deposition (17,18). Therefore, surface soil concentrations can be used to calculate a rough estimate of the amount of time which has passed since the soil was last inundated or wetted.

### Statistical Analyses

We performed a random forest analysis to determine if any of the measured environmental and geochemical variables could be used to predict whether a soil would have no detectable microorganisms or detectable microorganisms. The factors used in our models were chosen from a total of 37 different measurements. Highly correlated variables, edaphic factors, and variables that were not measured on at least 80% of the 204 samples were not included in the analysis. The final variables used for analysis included elevation, the concentration of certain water-soluble salts (NO_3_^-^, Cl^-^, ClO_3_^-^, ClO_4_^-^) leached at a 1:5 soil: water ratio, and a sum of total salt, total anions, and total cations calculated from these leached results. Cation and anion concentrations were log transformed before analysis. To determine what environmental variables might best predict which soils were uninhabited, we used the R package ‘rfPermute’ and performed a random forest analysis with 100 trees and 3 variables tried at each split to identify the most important predictors.

## Acknowledgments

We thank Jessica Henley, Matthew Gebert, Savanna Pierce, Brett Davidheiser-Kroll, Katie Snell, Sebastian Kopf, Marci Shaver-Adams, Natasha Griffin, Thomas Powers, Alyssa Pike, and Kevin Dickerson and Daniel Gilbert for their help with the laboratory analyses, Cecilia Milano de Tomasei for help with permits, stores, sample shipment, and Marci shaver-Adams and Geoff Schellens for help with sample collection and field safety support. We also thank the pilots and technicians of PHI helicopters, and the Shackleton Glacier camp staff for supporting our field campaign. Craig Cary provided feedback on several aspects of the project. This work was supported by grants from the U.S. National Science Foundation (ANT 1341629 to B.A., N.F., W.L., and D.W. and OPP 1637708 to B.A) with additional support provided to N.D. from University Colorado Department of Ecology and Evolutionary Biology.

## Data and materials availability

All data used in this study are available in the main text or the supplementary material with the raw sequence data from the cultivation-independent marker gene sequencing publicly available in GenBank (accession number xxxxx).

