## Supplemental Information for "Exploring the boundaries of microbial habitability in soil"

**This PDF file includes:**

Figures S1 to S5
Tables S1 to S4
SI References

**Other supplementary materials for this manuscript include the following:**

Datasets S1 to S3

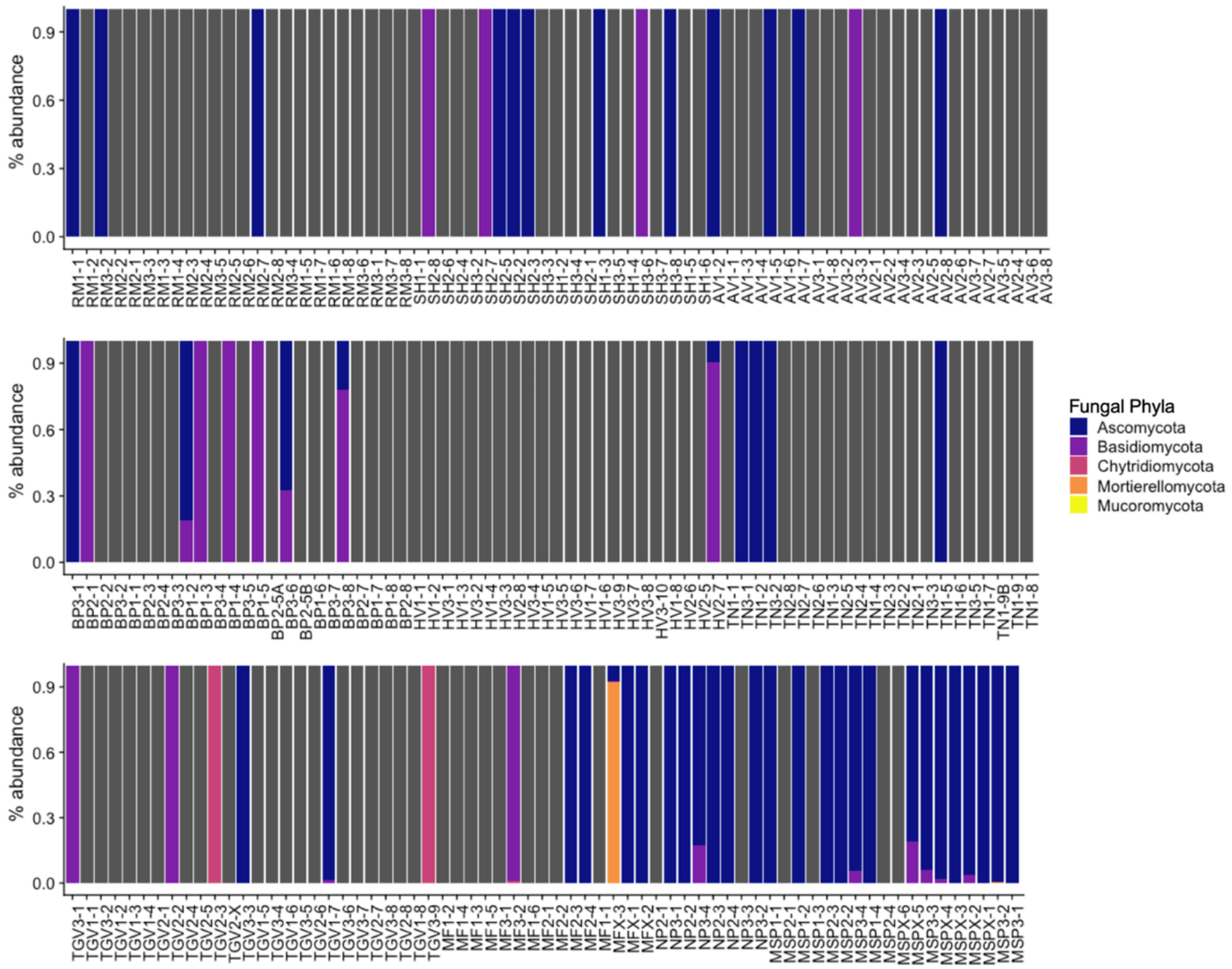


**Figure S1.** The relative abundances of fungal taxa identified using cultivation independent ITS gene sequencing in all 204 samples collected from across the Shackleton Glacier Valley. Samples are grouped by feature and organized from highest elevation site to lowest elevation site (top to bottom), with the higher elevation sites being further south. Grey bars represent samples with no amplifiable fungal DNA.

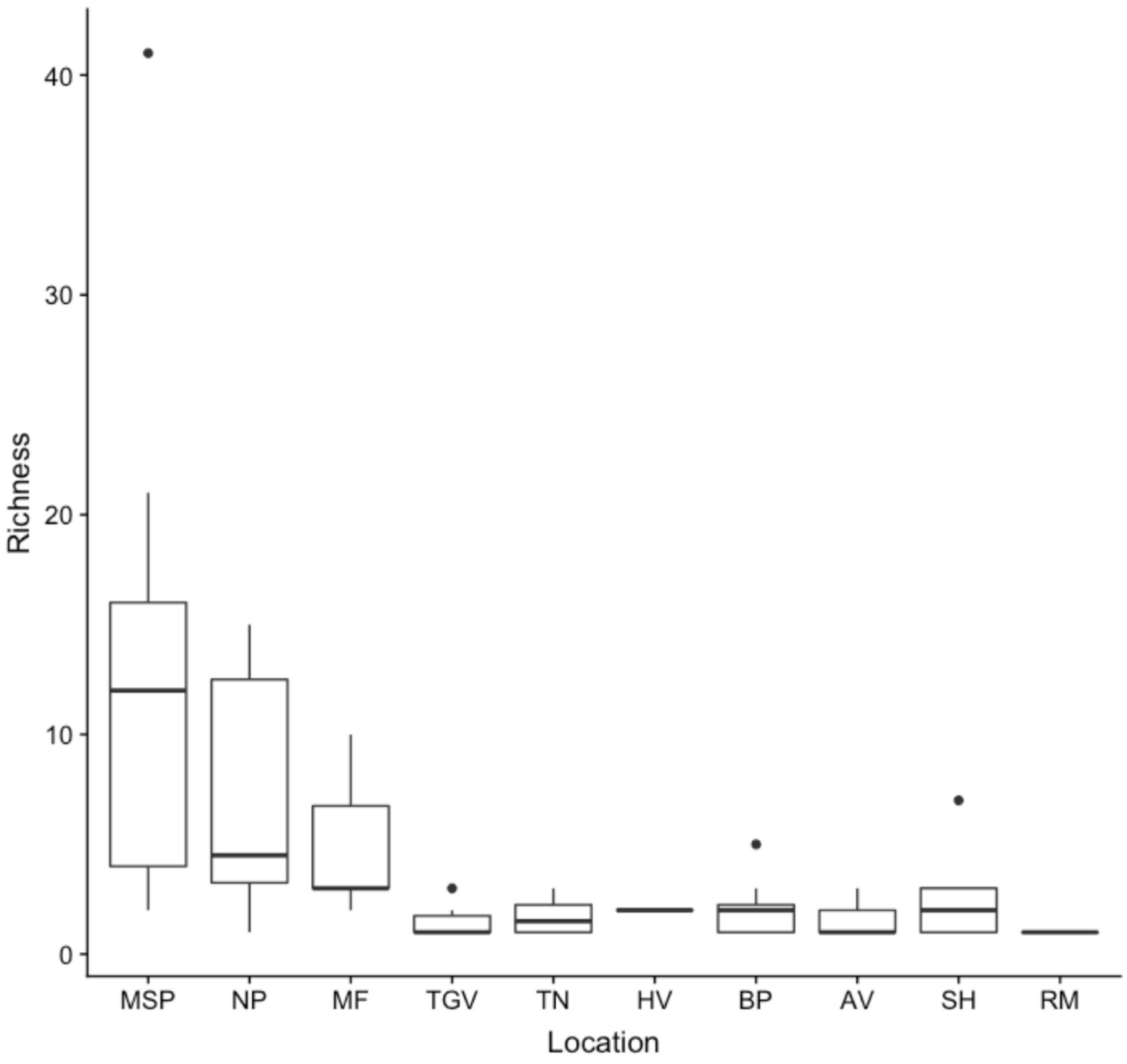


**Figure S2.** The richness of fungal communities across the 10 features of the Shackleton Glacier Valley. Fungal richness, number of unique fungal phylotypes per sample, was determined based on the cultivation independent ITS gene sequencing results. Samples were included if they had at least one read identifiable to a fungal phylum.

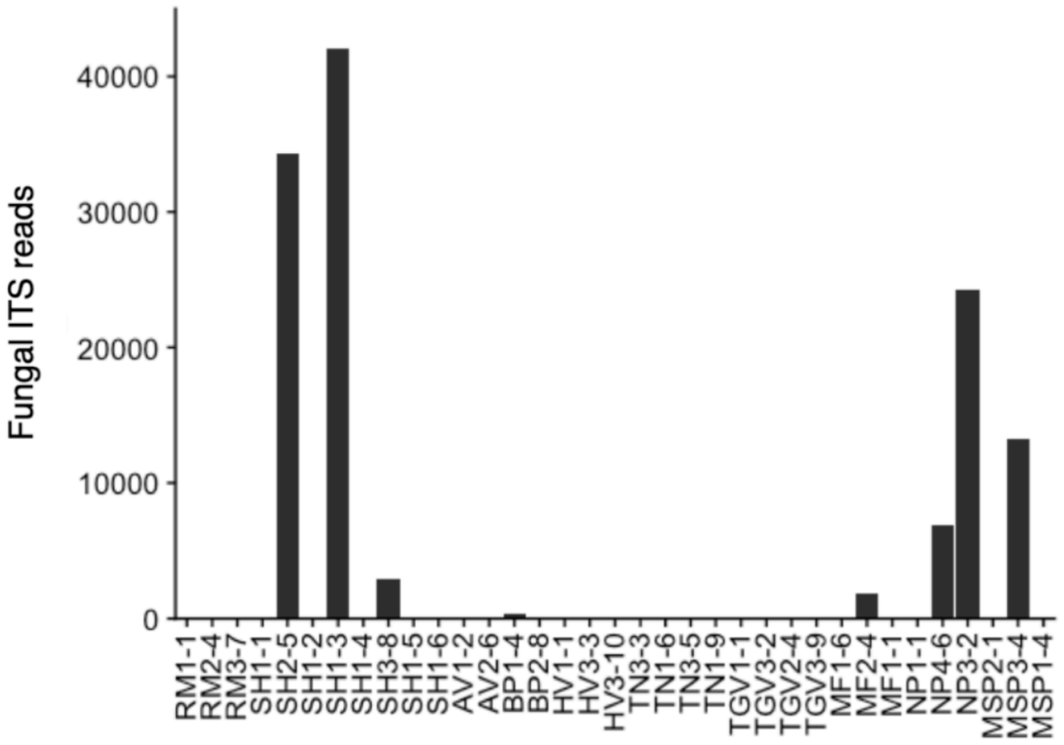


**Figure S3.** The number of fungal ITS reads that were classified at least to a fungal phylum for the same subset of 35 samples analyzed using the cultivation-dependent, metabolic, and ATP assays (see Figure 2A – 2D).

| **Sample** | **gDNA (ng/µl)** | **Amplified Product (ng/µl)** | **Well** |
|---|---|---|---|
| SH1-5 | 0.00 | 11.13 | B |
| SH1-5 + *E.coli* | 5.29 | 426.99 | C |
| SH1-6 | 0.00 | 15.91 | D |
| SH1-6 + *E.coli* | 8.49 | 407.58 | E |
| RM3-7 | 0.00 | 14.50 | F |
| RM3-7 + *E.coli* | 5.01 | 381.96 | G |
| TGV3-9 | 2.91 | 260.35 | H |
| TGV3-9 + *E.coli* | 17.28 | 370.79 | I |
| *E.coli* | 14.26 | 528.26 | J |
| Blank (Extraction) | 0.00 | 13.95 | K |
| Blank (PCR) | n/a | 15.95 | L |

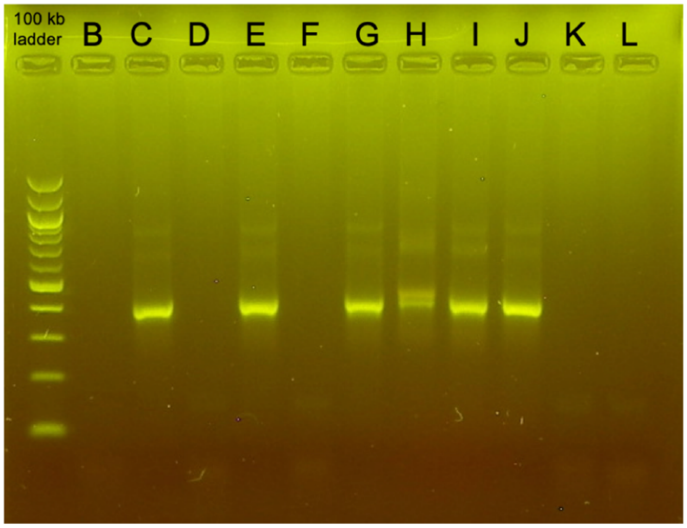


**Figure S4.** The results of the PCR inhibition test. The PCR inhibition test used three samples that were uninhabited (SH1-5, SH1-6, RM3-7) and one sample (TGV 3-9) that contained identifiable DNA based on the 16s rRNA sequencing. When the soils were spiked with *E. coli* cells, there was no evidence that DNA extractions or PCR amplifications were inhibited and thus the failure to detect amplifiable bacterial DNA in the 3 'uninhabited' soils is due to the lack of sufficient bacterial DNA in those samples and not methodological issue.

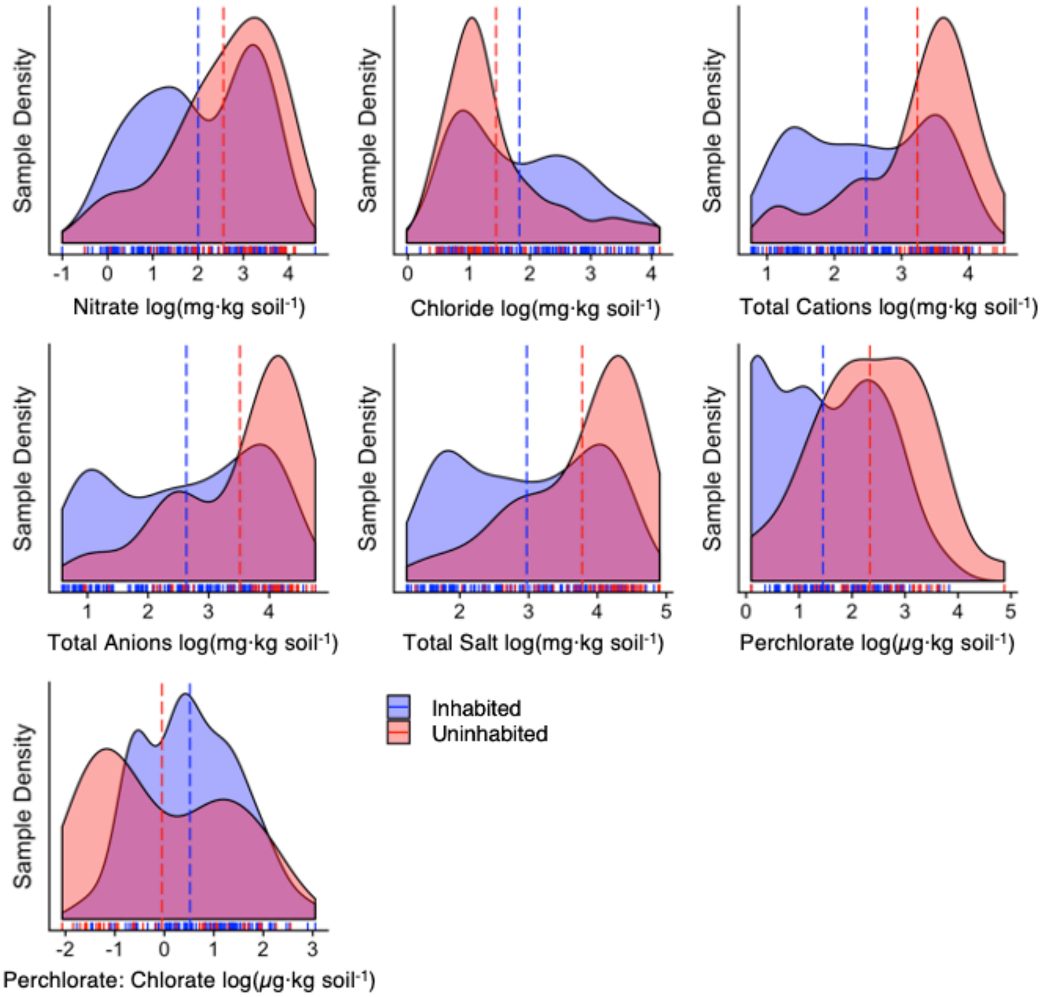


**Figure S5.** The distribution of the geochemical and environmental variables used in the random forest model that were not significant predictors of a soil's habitability. Y axis shows sample density of each habitability group with the number of samples (n) for each dataset ranging from 162-204. The area under each curve is equal to 1. Concentrations of all salt ions (μg·kg dry soil$^{-1}$, or mg·kg dry soil$^{-1}$) have been log transformed. Dashed lines indicate the average for each of the habitability groups. The model results for each of the different variables are indicated in Table S4.

**Table S1.** The growth media and growth conditions that were used for the culturing experiment.

| Media | pH | Temperature (°C) |
|---|---|---|
| Trypticase Soy Agar (TSA 24)[54] | 7.0 | 24 |
| Trypticase Soy Agar (TSA 4)[54] | 7.0 | 4 |
| M3 Acetate Agar (M3)[54] | 6.8 | 4 |
| Antarctic Bacterial Medium (ABM)[55] | 7.0 | 4 |
| Modified Nutrient Broth (MNB)[56] | 6.8 | 4 |
| Luria Bertani Media (LB)[54] | 7.0 | 4 |
| Malt Extract Agar (MEA)[54] | 5.5 | 4 |
| Photoautotrophic media (PA)[57] | 7.0 | 24 |
| Photoheterotrophic media (PH)[57] | 7.0 | 24 |
| VNSS agar (VNSS)[58] | 7.0 | 4 |
| V agar (V)[58] | 7.0 | 4 |
| R2A (pH_5)[59] | 5.0 | 4 |
| R2A (pH_7)[59] | 7.0 | 4 |
| R2A (pH_9)[59] | 9.0 | 4 |
| R2A+5% NaOH (Salt_5)[59] | 7.0 | 4 |
| R2A+10% NaOH (Salt_10)[59] | 7.0 | 4 |
| Potato Glucose Agar (PGA)[54] | 5.6 | 4 |

**Table S2.** The 35 samples used to confirm the results from the culture-independent genetic sequencing, their locations within the Shackleton Glacier Valley (see Methods).

| Sample | Site | Elevation (m) | Distance from Coast (km) | Relative age of last wetting (yrs) |
|---|---|---|---|---|
| AV1-2 | Augustana Valley | 1492 | 72 | 3.61E+03 |
| AV2-6 | Augustana Valley | 1376 | 72 | 4.06E+03 |
| BP1-4 | Bennett Platform | 1329 | 82 | 4.47E+03 |
| BP2-8 | Bennett Platform | 1222 | 82 | 3.47E+01 |
| MF1-1 | Mount Franke | 409 | 9 | 6.43E+01 |
| MF1-6 | Mount Franke | 484 | 9 | 3.47E+01 |
| MF2-4 | Mount Franke | 424 | 9 | 9.72E+00 |
| HV1-1 | Heekin Valley | 1660 | 63 | 8.68E+03 |
| HV3-3 | Heekin Valley | 1140 | 63 | 4.75E+03 |
| HV3-10 | Heekin Valley | 1030 | 63 | 5.86E+02 |
| MSP1-4 | Mount Speed | 188 | 0 | 3.47E+01 |
| MSP2-1 | Mount Speed | 270 | 0 | 3.47E+01 |
| MSP3-4 | Mount Speed | 193 | 0 | 3.47E+01 |
| NP1-1 | Nielsen Peak | | 0 | 3.47E+01 |
| NP3-2 | Nielsen Peak | 645 | 0 | 6.94E+00 |
| NP4-6 | Nielsen Peak | | 0 | 3.47E+01 |
| RM1-1 | Roberts Massif | 1801 | 120 | 1.19E+04 |
| RM2-4 | Roberts Massif | 1760 | 120 | 2.58E+02 |
| RM3-7 | Roberts Massif | 1688 | 120 | 6.44E+02 |
| SH1-1 | Schroder Hill | 2221 | 94 | |
| SH1-2 | Schroder Hill | 2123 | 94 | |
| SH1-3 | Schroder Hill | 2098 | 94 | 7.78E+04 |
| SH1-4 | Schroder Hill | 2091 | 94 | 1.12E+05 |
| SH1-5 | Schroder Hill | 2045 | 94 | 6.94E+04 |
| SH1-6 | Schroder Hill | 2039 | 94 | 5.53E+04 |
| SH2-5 | Schroder Hill | 2131 | 94 | 1.88E+05 |
| SH3-8 | Schroder Hill | 2057 | 94 | 4.31E+03 |
| TGV1-1 | Thanksgiving Valley | 1298 | 45 | 2.27E+04 |
| TGV2-4 | Thanksgiving Valley | 1086 | 45 | 2.38E+04 |
| TGV3-9 | Thanksgiving Valley | 911 | 45 | |
| TN1-6 | Taylor Nunatak | 955 | 45 | 2.44E+04 |
| TN1-9 | Taylor Nunatak | 883 | 45 | 1.72E+02 |
| TN3-3 | Taylor Nunatak | 1023 | 45 | 2.65E+04 |

**Table S3.** The results of the $^{13}C$ metabolism assay. The table shows the average % $^{13}C$ (+/- 1 standard deviation) recorded from the $^{13}CO2$ produced by the samples amended with glucose. 3 samples were classified as having no microbial activity detected, while 32 were classified as having microbial activity detected.

| Treatment Group | Unautoclaved $^{13}C$ ‰ | Autoclaved $^{13}C$ ‰ |
|---|---|---|
| Microbial activity detected | 9.90 +/- 9.65 | 1.15 +/- 0.09 |
| No microbial activity detected | 1.11 +/- 0.01 | 1.11 +/- 0.01 |
| Positive control | 15.91 +/- 10.01 | 1.10 +/- 0.01 |
| Blanks | 1.10 +/- 0.002 | |

**Table S4.** The importance of the predictive variables of the random forest in describing whether a soil had detectable microbial DNA or not. Salt ion concentrations (µg·kg dry soil$^{-1}$ or mg·kg dry soil$^{-1}$) were log-transformed before analysis. The importance of each variable is indicated by the % change of the mean standard error that is caused by the exclusion of that variable, and the significance (pval) of the estimation. This model explained 25.32% of the variance, with a mean of squared residuals of 0.130.

| Variable | % of MSE explained | p Val |
|---|---|---|
| Elevation | 7.85 | 0.02 |
| Chlorate | 7.66 | 0.01 |
| $ClO_4^-$: $ClO_3^-$: ratio | 4.32 | 0.06 |
| Total Salt | 4.02 | 0.44 |
| Total Cations | 3.50 | 0.45 |
| Perchlorate | 3.23 | 0.29 |
| $Cl^-$ | 2.40 | 0.45 |
| $NO_3^-$ | 2.36 | 0.77 |
| Total Anions | 0.46 | 0.98 |

**Data S1. (Separate File, Habitability_TaxTable_Bacteria.xlsx**)

The composition of all of the bacterial and archaeal taxa identified in the culture-independent 16s rRNA gene sequencing study for each of the 204 samples used. Taxonomic levels that are highlighted in this table are kingdom, phylum and family. Relative abundance of each taxonomic group listed is represented as % of total reads for that sample. Samples that were identified as containing no reliably detected prokaryotic DNA (see methods) show an abundance of 0% across all taxonomic groups.

**Data S2. (Separate File, Habitability_TaxTable_Fungi.xlsx**)

The composition of all of the fungal taxa identified in the culture independent ITS gene sequencing study for each of the 204 samples used. Taxonomic levels that are highlighted in this table are phylum and family. Relative abundance of each taxonomic group listed is represented as % of total reads for that sample. Samples that were identified as containing no reliably detected fungal DNA show an abundance of 0% across all taxonomic groups.

**Data S3 (Separate File, Habitability_ConfirmatoryTests_Results.xlsx)**

The results of the confirmatory tests for microbial habitability and activity (culture-independent, culture dependent, $^{13}C$ glucose assay, ATP Assay), using the subset of 35 samples (Table S2). The data included in table is visually represented in Figure 2 and includes the results of experimental blanks and positive controls.
